# The break in the cycle: Inositol pyrophosphate fluxomics disentangled via mathematical modelling

**DOI:** 10.1101/2025.03.27.645712

**Authors:** Hermes Jacques, Kim Geun-Don, Liu Guizhen, Maria Giovanna De Leo, Mayer Andreas, Jessen Henning, Timmer Jens

## Abstract

This study investigates the metabolic pathways of inositol pyrophosphates (IPPs) in the yeast cell line ΔSPX and the human tumor cell line HCT116. Utilizing pulse-labelling experiments with ^18^O water and ordinary differential equation (ODE) models, we explore the synthesis and turnover of the highly phosphorylated IPP, 1,5-InsP_8_. Our findings challenge the notion that 1,5-InsP_8_ can be synthesized through distinct routes, revealing a linear reaction sequence in both systems. Employing model reduction via the profile likelihood method, we achieved statistically concise identifiability analysis that led to significant biological insights. In yeast, we determined that 1,5-InsP_8_ production primarily occurs through the phosphorylation of 5-InsP_7_, with the pathway involving 1-InsP_7_ deemed unnecessary as its removal did not compromise model accuracy. In HCT116 cells, 1,5-InsP_8_ synthesis is mainly driven by 1-InsP_7_, with variations observed across different experimental conditions. These results underscore the utility of model reduction in enhancing our understanding of metabolic pathways, challenging traditional views of IPP metabolism, and providing a framework for future investigations into the regulation and implications of linear IPP pathways in eukaryotic cells.

## 1 Introduction

All living organisms utilize signalling molecules to respond to external stimuli and maintain internal stability. Inositol pyrophosphates (IPPs) are a class of signalling molecules that is evolutionarily conserved in eukaryotic cells [Holub, 1986]. IPPs directly bind to a family of conserved SPX domains (named from <u>S</u>yg1, <u>P</u>ho81 and <u>X</u>pr1), inducing conformational changes and regulating their functions [Wild et al., 2016, Ried et al., 2021, Pipercevic et al., 2023, Liu et al., 2023, Lu et al., 2024, Yan et al., 2024]. In addition, IPPs can induce non-enzymatic pyrophosphorylation of proteins to control their activities, such as Casein Kinase 2 (CK2), AP3, and dynein [Saiardi et al., 2004, Bhandari et al., 2007, Azevedo et al., 2009, Wu et al., 2016, Chanduri et al., 2016, Lolla et al., 2021, Morgan et al., 2024]. The regulation of protein activities by IPPs regulates key physiological processes, such as energy metabolism, hormone response, and phosphate homeostasis [Chakraborty, 2018, Thota and Bhandari, 2015, Austin and Mayer, 2020].

IPPs occurring in cells contain one or two pyrophosphate groups attached to the inositol ring. Their unique properties are determined by the number and positioning of the pyrophosphate groups, which may confer different physiological roles [Shears, 2018, Lee et al., 2020, Qi et al., 2023]. IPPs are synthesized by several kinases using InsP_6_ as a backbone. Inositol hexakisphosphate kinases (IP6Ks in mammals; Kcs1 in yeast) add a phosphate group at the C5 position of the inositol ring, generating 5-InsP_7_ from InsP_6_ or 1,5-InsP_8_ from 1-InsP_7_ [Saiardi et al., 1999, Saiardi et al., 2001]. Diphosphoinositol pentakisphosphate kinases (PPIP5Ks in mammals; Vip1 in yeast) phosphylate at the C1 position of the inositol ring and synthesize 1-InsP_7_ from InsP_6_ or 1,5-InsP_8_ from 5-InsP_7_ [Mulugu et al., 2007, Fridy et al., 2007]. Further IPPs have been identified in plants and mammalian cells [Gaugler et al., 2022, Qiu et al., 2023a], but at present it is not clear what physiological roles they might play and how they are synthesized. Highly phosphorylated IPPs are converted back to InsP_6_ by several phosphatases. PPIP5Ks are bidirectional enzymes containing both kinase and phosphatase activities [Dollins et al., 2020, Wang et al., 2015]. Their phosphatase domain can remove the *β*-phosphate at the C1 position from 1-InsP_7_ or 1,5-InsP_8_. In mammals, diphosphoinositol polyphosphate phosphohydrolases (DIPPs) nonspecifically remove the *β*-phosphate group, leading to the conversion of IPPs back to InsP_6_ [Safrany et al., 1998]. In yeast, several different phosphatases are involved in the dephosphorylation of IPPs. Ddp1, a yeast homolog of mammalian DIPPs, preferentially removes the *β*-phosphate group at the C1 position, generating 5-InsP_7_ from 1,5-InsP_8_ or InsP_6_ from 1-InsP_7_ [Safrany et al., 1999, Kilari et al., 2013, Marquez-Monino et al., 2021]. Meanwhile, Siw14 remove the *β*-phosphate at the C5 position to create 1-InsP_7_ from 1,5-InsP_8_ or InsP_6_ from 5-InsP_7_ [Steidle et al., 2016, Benjamin et al., 2022, Qin et al., 2023].

The levels of IPPs dynamically change quantitatively and qualitatively in response to various cellular signals and environmental stimuli. For example, when yeast cells are starved for inorganic phosphate (P_i_), the amount of overall IPPs decrease rapidly, with 1,5-InsP_8_showing a faster and more pronounced decline than 1-InsP_7_ and 5-InsP_7_ [Chabert et al., 2023]. The decrease in 1,5-InsP_8_ triggers the phosphate-responsive signal transduction (PHO) pathway. This pathway induces transcription of genes that increase the capacity for scavenging P_i_ from the surrounding environment, and for liberating P_i_ from polyphosphates, nucleic acids, or lipids to enable internal P_i_ recycling. 1,5-InsP_8_ responds similarly to P_i_ starvation in mammalian cells and in plants, suggesting that its role in P_i_ signalling is conserved [Menniti et al., 1993, Dong et al., 2019, Li et al., 2020]. The amount of IPPs also changes during the cell cycle, perhaps as a consequence of fluctuating P_i_ utilization [Barker et al., 2004, Banfic et al., 2013, Bru et al., 2016, Neef and Kladde, 2003]. Changes of IPPs in turn affect the cell cycle progression, e.g., by controlling kinetochore formation [Kuenzel et al., 2022]. In addition, environmental stresses such as hyperosmotic stress [Pesesse et al., 2004], oxidative stress [Steidle et al., 2020, Eisenbeis et al., 2023], and heat stress [Choi et al., 2005] can affect IPPs levels.

1,5-InsP8 can be produced through two separate routes from IP6, via 5-InsP_7_ or 1-InsP_7_. It was proposed that the main route for synthesizing 1,5-InsP_8_ from InsP_6_ is through 5-InsP_7_, while the conversion of InsP_6_ to 1-InsP_7_ by PPIP5Ks is not efficient in vivo [Randall et al., 2020]. This model was derived from several lines of evidence: 1) The cellular concentration of InsP_6_ is markedly higher than that of other IPPs, but its turnover rate is significantly slower compared to that of other IPPs [Glennon and Shears, 1993], implying that a substantial amount of InsP_6_ is compartmentalized and remains unreactive. Perhaps through cation-mediated ion bridges with the membrane [Poyner et al., 1993]. 2) Several studies showed that PPIP5Ks prefer 5-InsP_7_ as a substrate over InsP_6_ [Weaver et al., 2013,Gu et al., 2017]. 3) The concentration of 1-InsP_7_ within the cell is very low, almost to the point of being undetectable under normal conditions [Gu et al., 2016,Chabert et al., 2023]. While substrate preferences of the IPP enzymes could be dissected through in vitro studies [Weaver et al., 2013, Gu et al., 2017], such information does not automatically predict the metabolite fluxes in vivo. Even if a preference for 5-InsP_7_ exists, this does not exclude InsP_6_ as a potential substrate. Therefore, it is required to analyse IPPs synthesis in vivo to understand the 1,5-InsP_8_ synthesis pathway. Here we used data from pulse-labelling experiments with ^18^O water to dissect the kinetics of IPP turnover and ordinary differential equation (ODE) models to reveal the relevant metabolite fluxes. In order to estimate the parameters of our ODE models, we used the modelling tool box *dMod* [Kaschek et al., 2019], which was developed in our group, to fit the ODE model to the pulse-labelling data. Furthermore we applied a method called profile likelihood [Raue et al., 2013, Kreutz et al., 2013] to perform a statistical analysis of the estimated parameters’ uncertainties. Based on this we reparameterized our models, i.e. adjusted the model structures, in a process called model reduction [Maiwald et al., 2016,Tönsing et al., 2018] and found that the analysed organisms, the yeast Saccharomyces cerevisiae and a human tumor cell-line (HCT116), synthesize 1,5-InsP_8_ through distinct routes. In both cases however, the statistically preferred version of their IPP metabolic routes is not a cycle at all but rather a linear reaction network.

## 2 Results

In this work we analysed two different biological organisms, baker’s yeast and HCT116. The data sets were first analysed using the same initial, in the following called “full”, mathematical model (Fig. 1). The data-sets analysed in this manuscript stem from a recently published approach, in which cellular IPs are rapidly pulse-labelled through ^18^O water [Kim et al., 2024, Qiu et al., 2023b]. Since it is expected, that the metabolic cycle has different kinetics between both organisms, the model was fitted and reduced for both organisms independently. However the applied methods and strategies were identical in all analyses to ensure statistically sound results. For every iteration a multi-start fit approach was chosen, i.e. multiple fits with individual and randomly sampled initial parameter vectors were performed. To obtain statistically sound and comparable results, 200 multistarts were performed for each iteration. In the following, the model reduction steps leading to the final fully reduced model are described for both organisms independently. The mathematical modelling techniques employed in this work are explained in detail in the Materials and Methods section.

**Figure 1:**
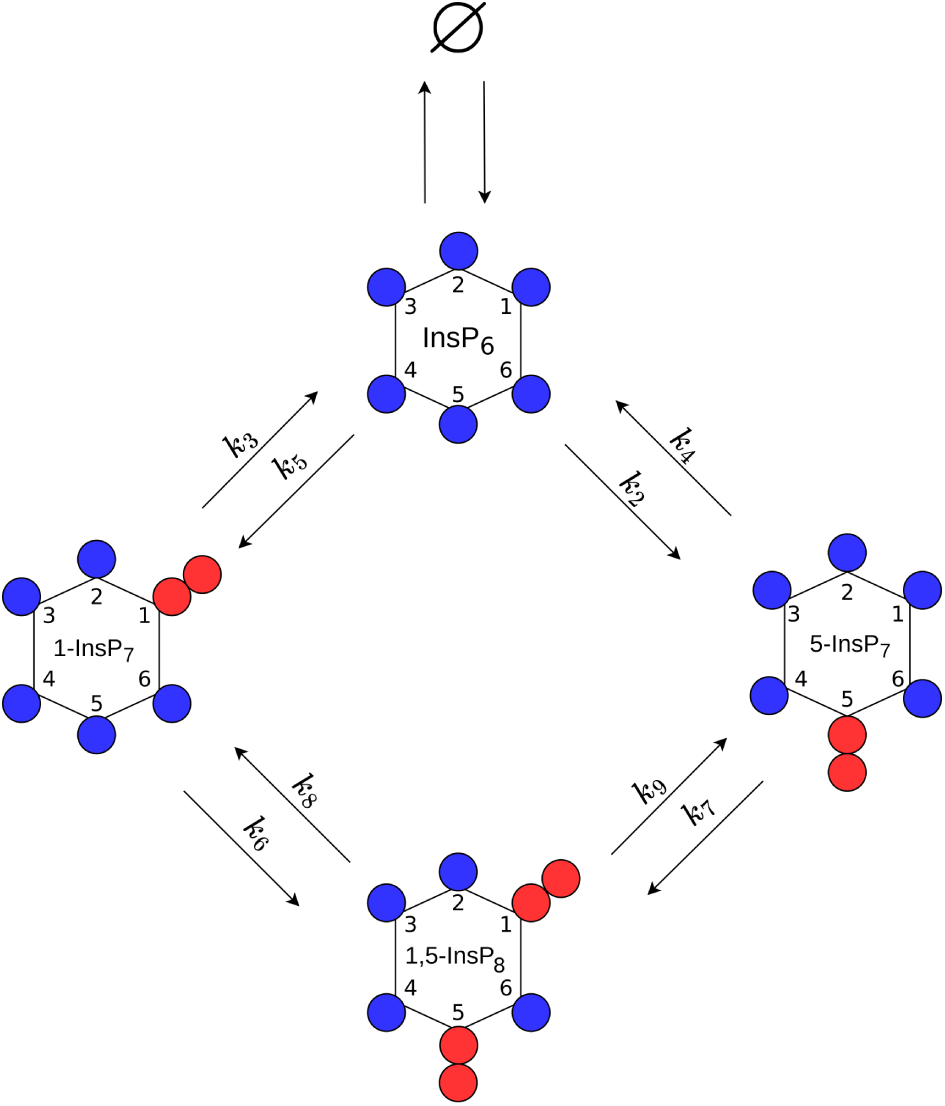
Model scheme of the cyclic production pathway of 1,5-InsP_8_ upon which the “full” ODE-model is based.

### 2.1 Yeast

In the yeast dataset, the levels of 1-InsP_7_ were not shown because of their low abundancein the samples [Kim et al., 2024]. For that reason, we reinvestigated the ^18^O labelling of IPPs under the high P_i_ conditions in the cytosol, which favours the accumulation of all IPP species [Chabert et al., 2023]. To this end, we overexpressed the truncated form of the low-affinity phosphate transporter Pho90 (Pho^ΔSPX^), which lacks the restraining SPX domain and allows enhanced P_i_ import into the cells [Hürlimann et al., 2009]. The cells were first grown in low phosphate medium (0.2 mM) and then transferred to high phosphate medium (50 mM) with 50% ^18^O water to generate P_i_ influx and enhanced IPP synthesis. For further details on the generation of the data set, we refer to the Materials and Methods section. In a first step, the yeast data-set was analysed using the “full” ODE model, which yielded a set of best-fit parameters which adequately described the dynamics observed in the data and yielded a Bayesian information criterion value (BIC) of -1011,9. The data as well as the time-courses obtained via the mathematical model with best-fit parameters are shown in panel A of Fig. 2. In panel B, the likelihood values of the 50 best fits out of the 200 multi-starts are shown sorted by the lowest likelihood. Such a plot, called waterfall plot, indicates that the best fit found in the analysis is found by eight different initial parameter configurations, providing evidence that the found optimum is in fact a global and not just a local one [Raue et al., 2013]. In a next step, parameter profiles were calculated, as shown in Fig. 2 C. This resulted in six model reactions where the rate parameters *k_i_* and the Michaelis-Menten (MM) constant *K_Mi_* are both not identifiable, namely *i* ∈ {2, 3, 5, 6, 7, 8}. A deviation from the profiled parameters best-fit value can thus be compensated by the re-optimization of other parameter values resulting in the same likelihood values, therefore being called coupled to or dependent on the compensating parameters. The dependencies of the nonidentifiable *k_i_*s and *K_M_* s are graphically displayed in Fig. 3 where the two strongest dependencies are displayed as paths in red and blue respectively and the dependencies of the remaining model parameters are shown in gray. In most panels, only the strongest dependency in red is visible since that parameter only strongly depends on one other parameter and the gray lines are thus overlapping the blue line. The parameter profile of interest is again plotted above its respective paths. Expect for *k*_6_ and *k*_8_, the parameters solely depend on the *K_M_* or *k* values of their respective reactions. Furthermore, if the profiled parameter value is increased the respective counterpart also increases in order to compensate for the change. Combining this observation with the fact that the *K_M_* values of interest have profiles with confidence intervals open to +∞ we see they can take very large values without worsening the likelihood. This supports reducing these reactions to mass-action reactions. Such mass-action reactions occur in a regime of Michaelis-Menten (MM) reactions where the *K_M_* value is much larger than the available substrate concentration. For *k*_6_ and *k*_8_ this reduction is also valid, since their *K_M_*values, which are the parameters that are being reduced, only depend on their respective *k* values. However, we do not expect *k*_6_ and *k*_8_ to become identifiable via this step, due to their strong dependencies upon each other, as seen in Fig. 3.

**Figure 2:**
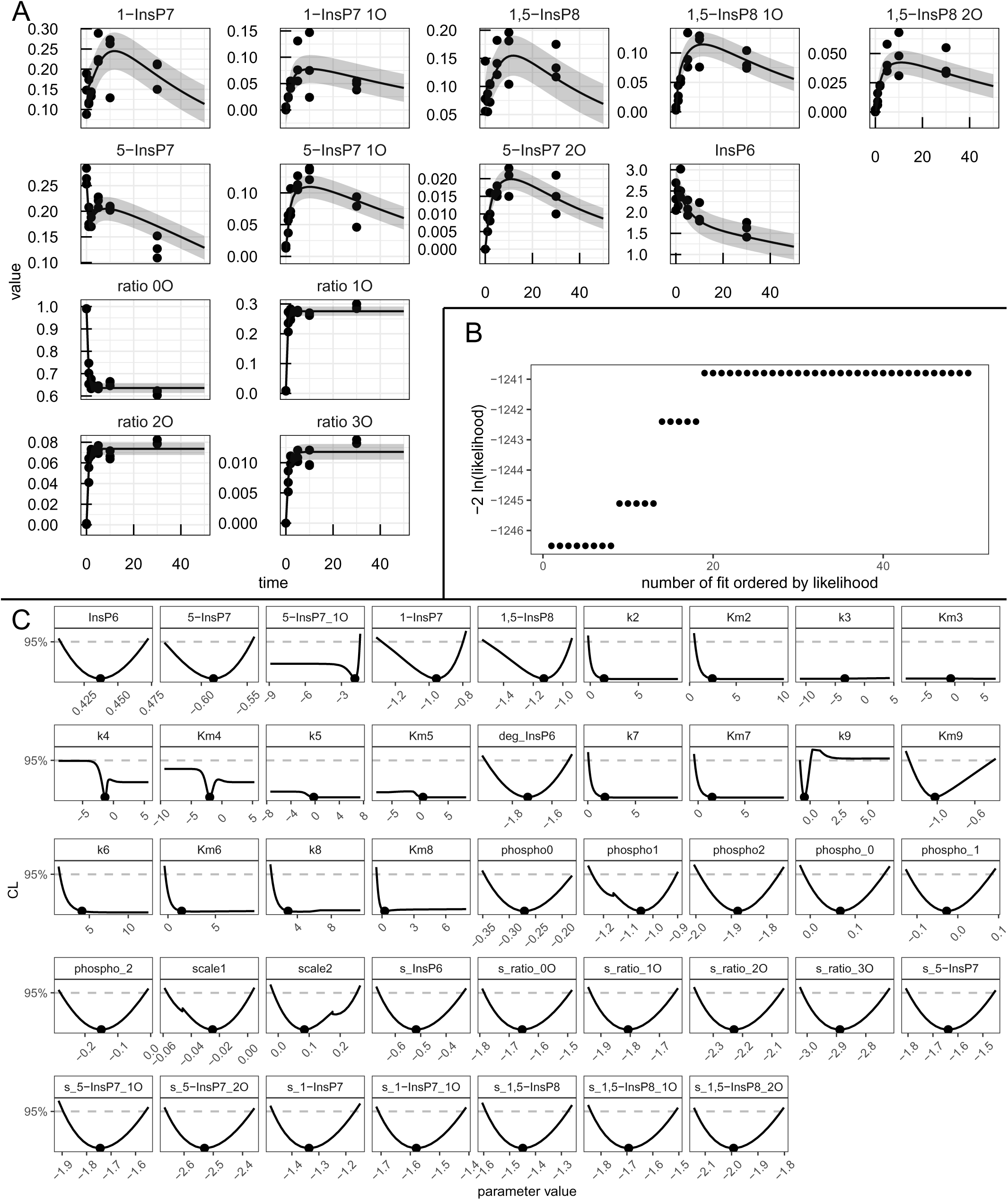
Analysis of the yeast data-set with the “full” mathematical model. (A) Fit of the mathematical model with the best-fit parameters (line) to the experimental data (points). The values are given in *µ*M except for the four species in the lower left of the panel which represent ratios of 1,2,3-labelled to 0-unlabelled ATP *γ*-phosphate. (B) Likelihood values of the 50 best multi-start fits, ordered by lowest likelihood value. (C) Profiles of the best-fit parameters of the full model (line). Best-fit parameter value are represented as points. Parameter values (x-axis) are displayed on log10 scale.

**Figure 3:**
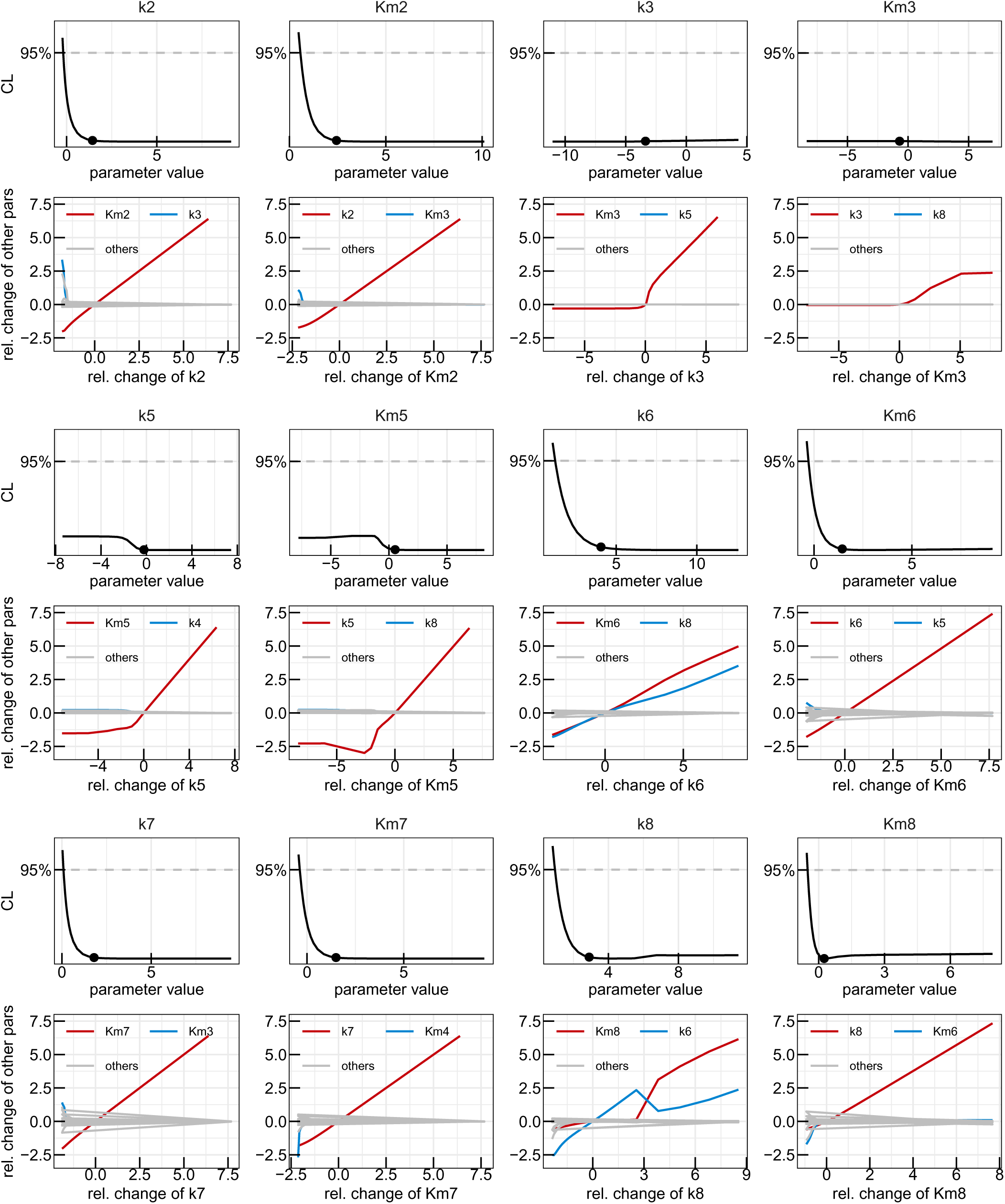
First reduction step in the yeast model. Profiles of the parameters considered for the first model reduction step in the yeast metabolic cycle. Best-fit parameter value are represented as points. Parameter values (x-axis) are displayed on log10 scale. Below each profile, the paths visualizing the dependencies on the other model parameters are plotted, with the strongest dependency in red, next strongest in blue and the remaining dependencies in gray. Relative change of the profiled parameter (x-axis) and relative change of the other parameters along the profile (y-axis) are displayed on log10 scale.

A subsequent multi-start fit of the so reduced model to the same data set showed no visible changes in goodness-of-fit, while BIC improved to -1045,2. Examining the parameter profiles resulting from this refitting implied that also the reaction rates *k*_3_ and *k*_5_ could be set to zero, since their profiles are open towards -∞ on the log scale (or zero on the linear scale), as shown in Fig. 4 A. Furthermore, no significant dependencies for small values of these parameters are present, demonstrating that for values smaller then the best-fit one, other parameters do not need to be adjusted in order to keep the best likelihood value, justifying setting these reaction rates to zero on linear scale. The profiles of both *k*_8_ and *k*_6_ on the other hand are still open towards +∞ on the log scale, meaning both reactions could be set to arbitrarily high values without significantly changing the model’s capacity of describing the data. The remaining dependencies of the two parameters in Fig. 4 A indicate that they linearly dependent upon one-another signifying that one can introduce the parameter transformation 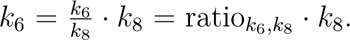 It is expected, that the ratio between the two rates becomes identifiable.

**Figure 4:**
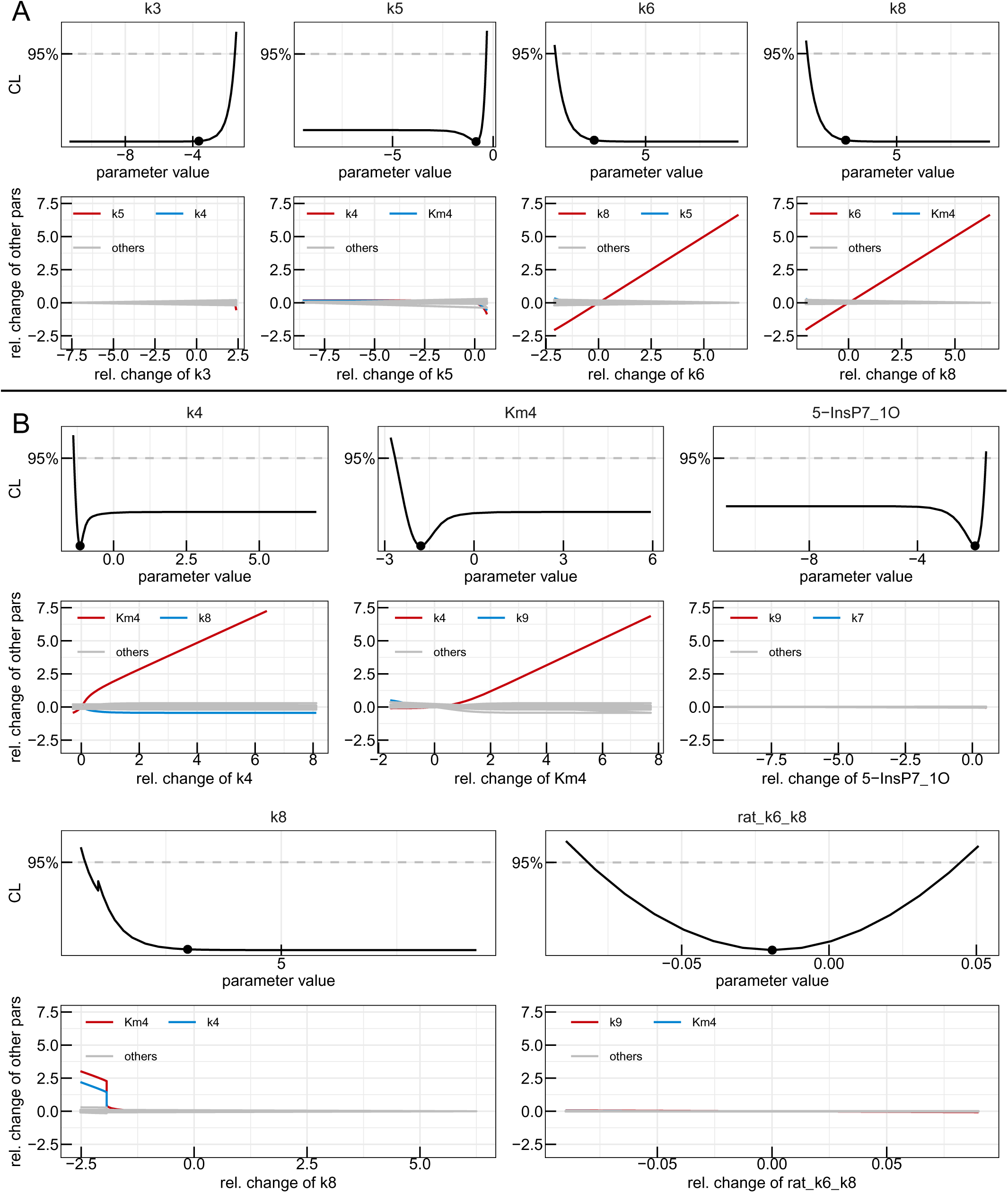
Second and third reduction steps in the yeast model. Profiles of the parameters considered for the second (A) and third (B) model reduction step in the yeast metabolic cycle. Best-fit parameter value are represented as points. Parameter values (x-axis) are displayed on log10 scale. Below each profile, the paths visualizing the dependencies on the other model parameters are plotted, with the strongest dependency in red, next strongest in blue and the remaining dependencies in gray. Relative change of the profiled parameter (x-axis) and relative change of the other parameters along the profile (y-axis) are displayed on log10 scale.

The now twice reduced model was once again fitted to the data, yielding an unchanged description of the data while resulting in a BIC of -1055,9. In a last reduction step, the reaction involving *k*_4_ and *K_M_*_4_ are reduced to mass action, following the same reasoning as for the other mass-action reduction steps. Additionally, the initial value for single labelled 5-InsP_7_ was also reduced to 0 following the reasoning of the previous reductions of *k*_3_ and *k*_5_. The profiles and dependencies of these three parameters are shown in Fig. 4 B, together with the identifiable profile of the ratio between *k*_6_ and *k*_8_ and the profile of *k*_8_ which, as expected, is still open towards +∞. Since the goodness-of-fit is not limited by large values of *k*_8_ its value was fixed to 10^5^ on the linear scale. The final, completely reduced model yielded profiles shown in Fig. 5 A and a BIC of -1069.5. It is completely identifiable as seen in the parabolic shapes of every single calculated profile. Note, that only the description of the reaction involving *k*_9_ requires the use of Michaelis-Menten kinetic since both *k*_9_ and *K_M_*_9_ are identifiable parameters. In panel B of Fig. 5 the waterfall plot of the 50 best multi-starts is plotted, where no step can be observed since all 50 fits ended up at the same best likelihood value, which indicates high trust in the employed numerical optimisations. Finally, in panel C, a scheme of the statistically favoured version of the reaction pathway is depicted, with red crosses marking the reactions which were reduced to 0. This scheme is not a cycle anymore, but rather a chain where production of 1-InsP_7_ can only be facilitated via dephosphorylation of 1,5-InsP_8_ rather than phosphorylation of IP6.

**Figure 5:**
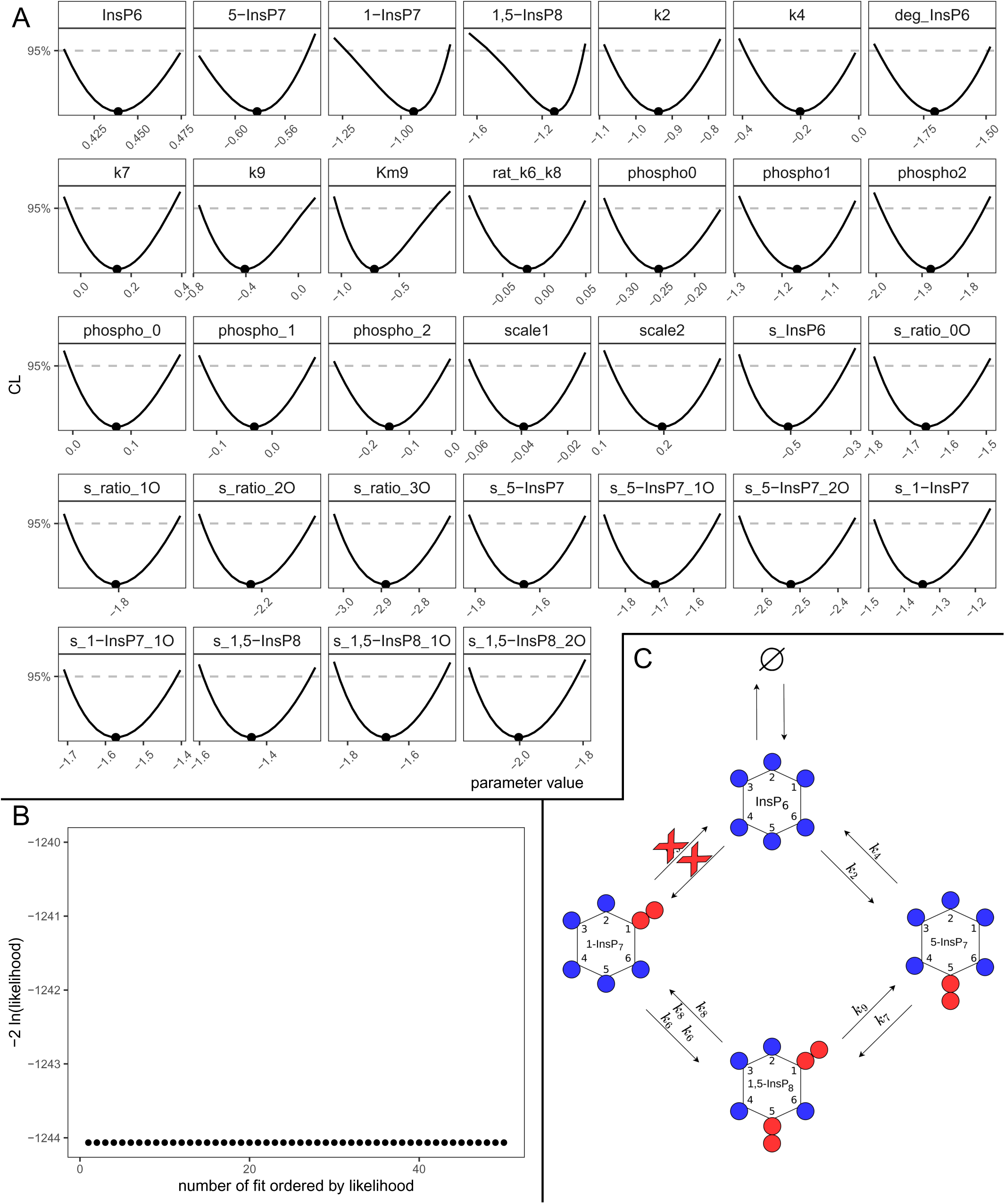
Fully reduced yeast model. (A) Profiles of the best-fit parameters of the reduced model (line). Best-fit parameter value are represented as points. Parameter values (x-axis) are displayed on log10 scale. (B) Likelihood values of the 50 best multi-start fits, ordered by lowest likelihood value. (C) Representation of the statistically favored transition scheme, displaying a chain-like pattern rather than a cycle.

### 2.2 HCT116

Human HCT116 cells were grown in normal phosphate medium and then transferred to normal phosphate medium with 50% ^18^O water. For further details on the generation of the data sets are given in Materials and Methods. The first fit with the “full” model yielded again an adequate description of the dynamics present in the data with a BIC of 164,6. The HCT116 data set together with the “full” model fit is depicted in Fig. 6A. A waterfall plot displaying the sorted likelihood values of the 50 best fits is shown in Fig. 6B, where eleven different initial configurations ended up in the same global optimum. Analysis of the model parameters profiles, displayed in Fig. 6C, shows that also in this case a number of parameters are unidentifiable, indicated by a non-parabolic profile. Thus model reduction is required. The parameters of interest in the first model reduction step together with their respective paths can be found in Fig. 7. The reactions involving *k*_2_, *k*_4_ and *k*_5_ are reduced to mass action following the reasoning established in Section 2.1. Furthermore the parameters *k*_7_ and *K_M_*_3_ are set to zero, eliminating the transition involving *k*_7_ from the model, while reducing the *k*_3_ reaction to a zero order reaction. In a zero order reaction, as explained in Section4.1, the reaction rate is independent of the substrate concentration.

**Figure 6:**
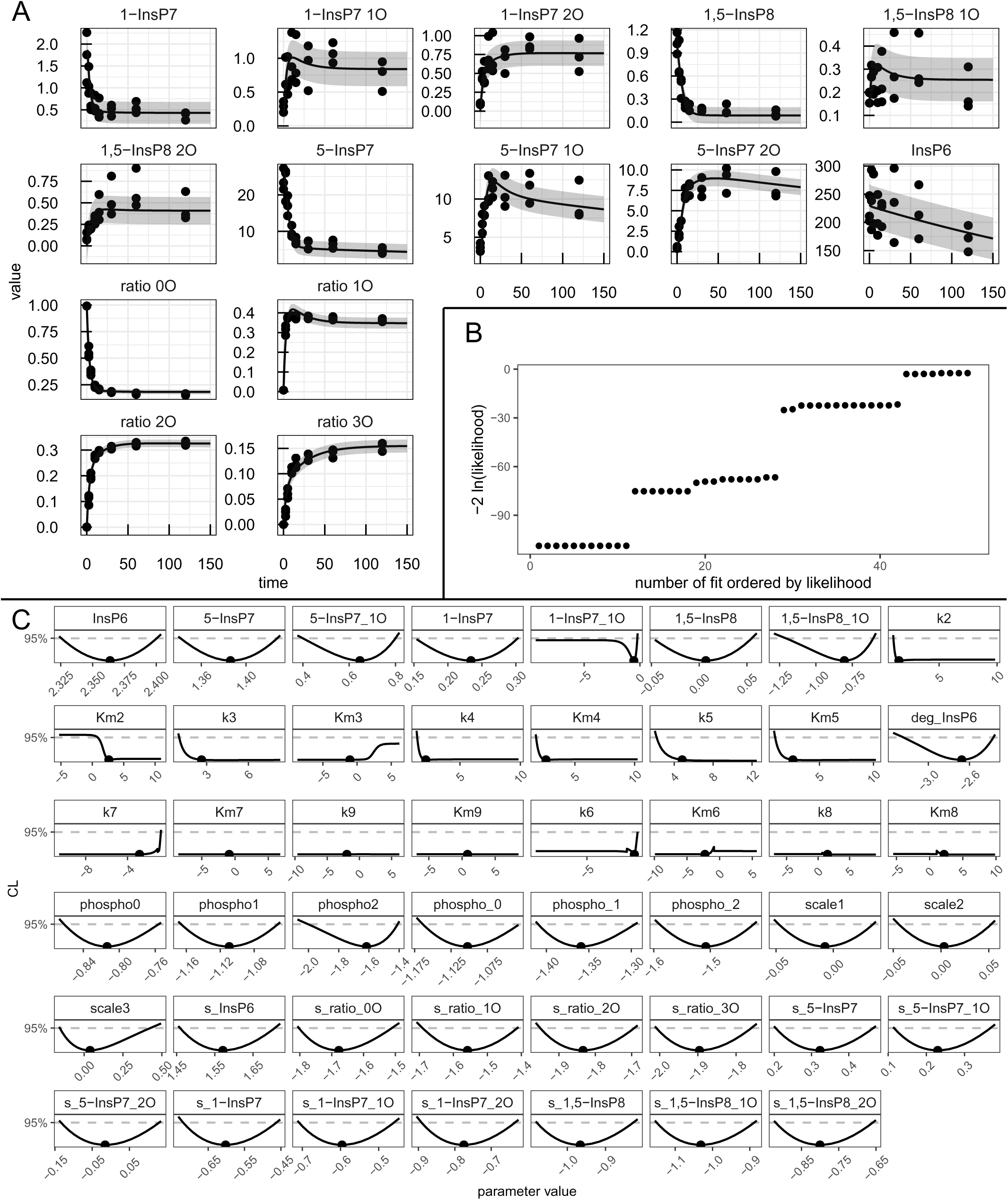
Analysis of the HCT116 data-set with the “full” mathematical model model. (A) Fit of the mathematical model with the best-fit parameters (line) to the experimental data (points). The values are given in *µ*M except for the four species in the lower left of the panel which represent ratios of 1,2,3-labelled to 0-unlabelled *γ*-phosphate. (B) Likelihood values of the 50 best multi-start fits, ordered by lowest likelihood value. (C) Profiles of the best-fit parameters of the full model (line). Best-fit parameter value are represented as points. Parameter values (x-axis) are displayed on log10 scale.

**Figure 7:**
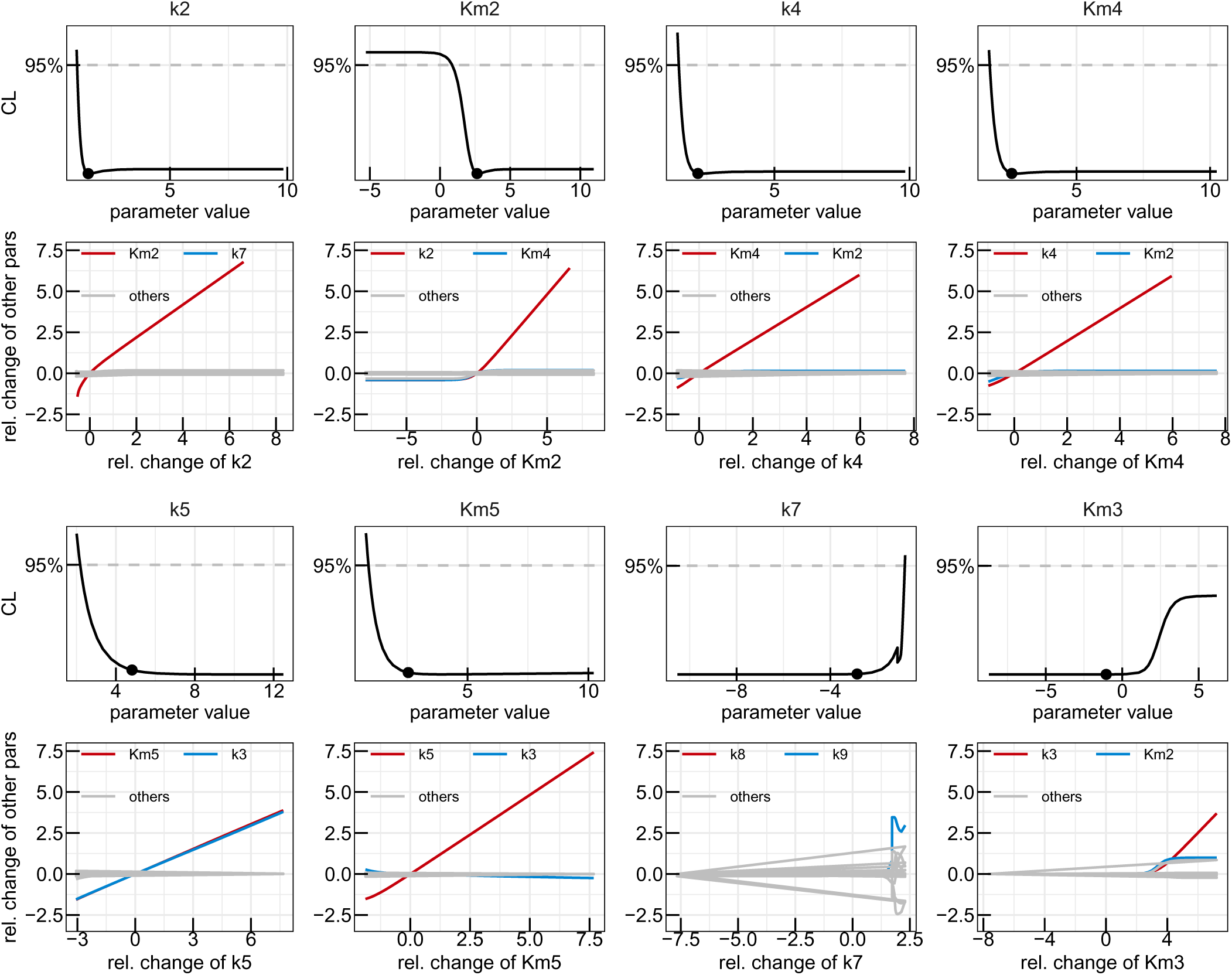
First reduction step in the HCT116 model. Profiles of the parameters considered for the first model reduction step in the yeast metabolic cycle. Best-fit parameter value are represented as points. Parameter values (x-axis) are displayed on log10 scale. Below each profile, the paths visualising the dependencies on the other model parameters are plotted, with the strongest dependency in red, next strongest in blue and the remaining dependencies in grey. Relative change of the profiled parameter (x-axis) and relative change of the other parameters along the profile (y-axis) are displayed on log10 scale.

Refitting the HCT16 data-set with the reduced model yielded no visible changes in data description but resulted in an improved BIC of 128,9. In Fig. 8 A, the reduction to mass-action reactions can be appreciated for the *k*_8_ and *k*_9_ transitions. Even though, *K_M_*_8_ shows a strong dependency on *k*_9_ for small values, for large values *K_M_*_8_ only depends linearly on its corresponding reaction rate, legitimising this reduction. Similar to *k*_6_ and *k*_8_ in Section 2.1, here *k*_3_ and *k*_5_ depend linearly on each other while having profiles which are open towards +∞, demanding the parameter transformation 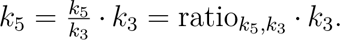 Lastly, the reaction involving *k*_6_ is reduced to a zero order reaction following the example of *k*_3_.

**Figure 8:**
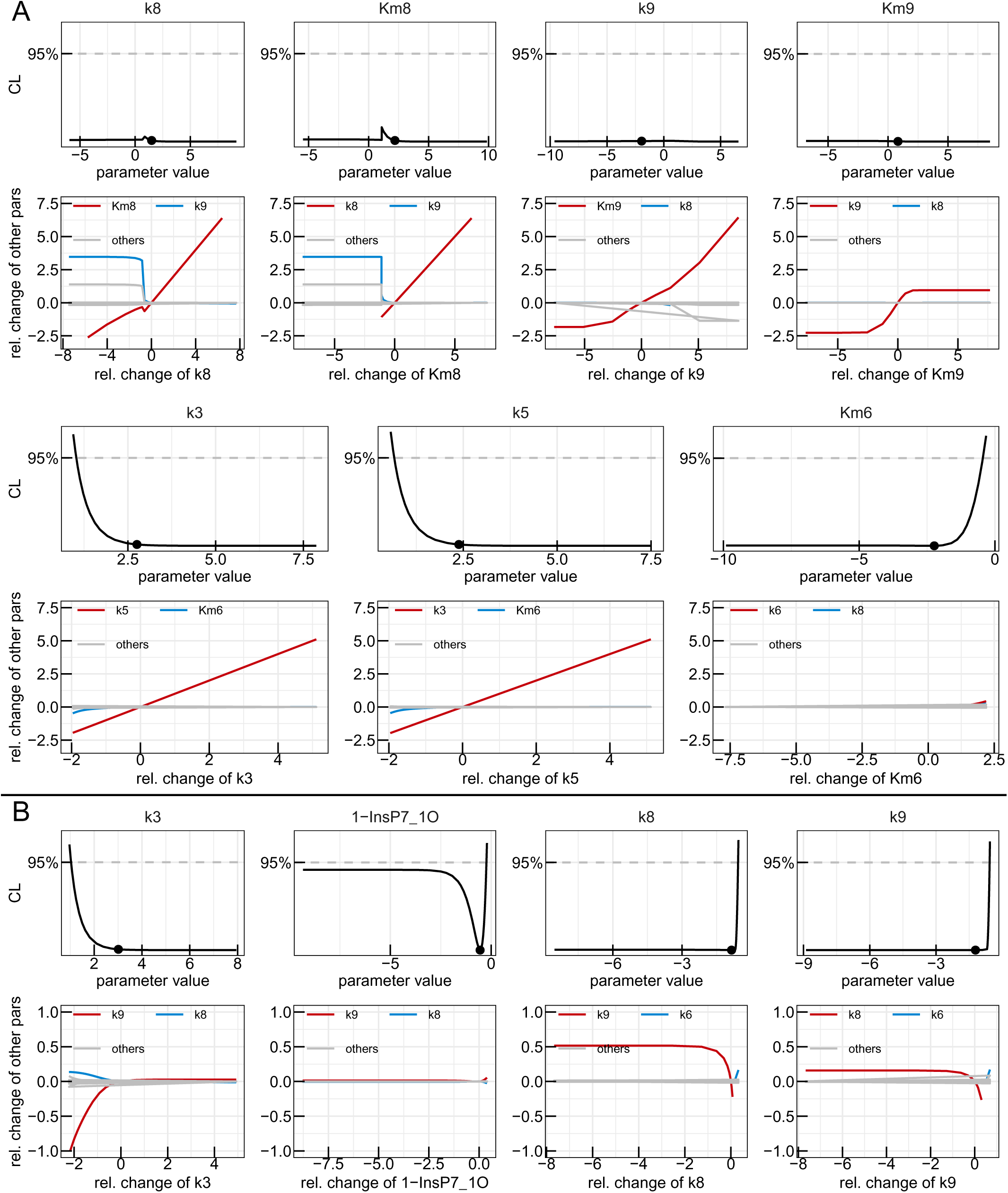
Second and third reduction steps in the HCT116 model. Profiles of the parameters considered for the second (A) and third (B) model reduction step in the yeast metabolic cycle. Best-fit parameter value are represented as points. Parameter values (x-axis) are displayed on log10 scale. Below each profile, the paths visualising the dependencies on the other model parameters are plotted, with the strongest dependency in red, next strongest in blue and the remaining dependencies in grey. Relative change of the profiled parameter (x-axis) and relative change of the other parameters along the profile (y-axis) are displayed on log10 scale.

This version of the model can describe the data without a visible change in goodness-of-fit, but yielded a BIC of 111,1. In a last reduction step, the parameters shown in Fig. 8 B are considered. *k*_3_ is set to 10^5^, while the initial value of single labelled 1-InsP_7_ is set to 0. When choosing between *k*_8_ and *k*_9_, the latter is set to zero, since it shows negligible dependence on *k*_8_ compared to vice versa.

The final fit of the fully reduced model to the data generated a BIC of 96,94. The fully identifiable model parameters are shown in Fig. 9A. In this panel, the identifiable ratio between *k*_5_ and *k*_3_ can be appreciated as well as the fact that all of the remaining reactions could be reduced to mass action while still describing the data accurately. In the final waterfall plot of the fully reduced model (Fig.9B 26 fits ended up in the global optimum. Thus also the IPP metabolic cycle of the HCT116 appears to be linear rather than cyclic (Fig. 9C), where 1,5-InsP_8_ is solely generated by phosphorylation of 1-InsP_7_.

**Figure 9:**
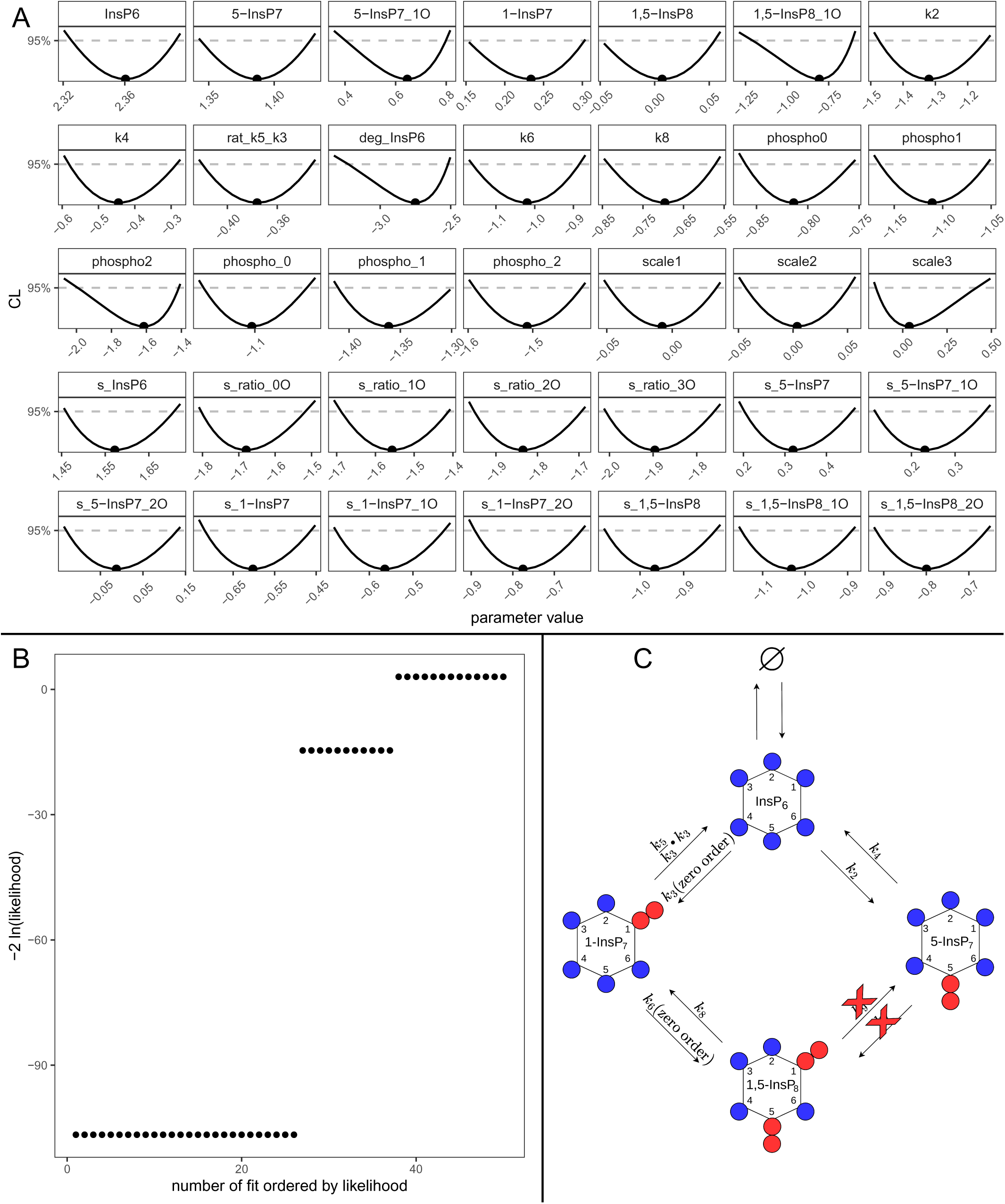
Fully reduced HCT116 model. (A) Profiles of the best-fit parameters of the reduced model (line). Best-fit parameter value are represented as points. Parameter values (x-axis) are displayed on log10 scale. (B) Likelihood values of the 50 best multi-start fits, ordered by lowest likelihood value. (C) Representation of the statistically favoured transition scheme, displaying a chain-like pattern rather than a cycle.

For HCT116, a second data set was generated and analysed, where the experimental conditions of the yeast experiment were copied, regarding the transition from low to high phosphate. The fits and the model reduction can be found in the supplement (Figs. 10-13). The results are consistent with the analysis of the first HCT116 dataset in the sense that also in this reduced model, the transitions between 5-InsP7 and 1,5-InsP8 are removed. The transition between 1-InsP7 and InsP_6_ is also removed.

## 3 Discussion

In this study, we investigated the metabolic pathways of inositol pyrophosphates (IPPs) in two different biological systems: yeast and the human tumour cell line HCT116. By utilising pulse-labelling experiments with ^18^O water and ordinary differential equation (ODE) models, we explored the synthesis and turnover of IPPs, aiming to clarify the pathways involved in the production of the highly phosphorylated IPP, 1,5-InsP_8_. Previous studies had proposed that 1,5-InsP_8_ could be synthesised via two distinct routes: either through 5-InsP_7_ or 1-InsP_7_. Our findings challenge this notion, suggesting that the metabolic networks in both yeast and HCT116 cells do not form a cyclic pathway but follow a linear reaction sequence.

We first applied a comprehensive (”full”) ODE model to describe the dynamics of IPP metabolism in yeast and HCT116 cells. The initial fitting results revealed that while the model could adequately capture the data, several parameters were not identifiable, indicating redundancies and dependencies that made it challenging to precisely estimate certain reaction rates and Michaelis-Menten constants. To address these issues, we performed a systematic model reduction process, heavily based on the profile-likelihood method. This process involved simplifying the model by reducing certain reactions to mass-action kinetics or even setting specific reaction rates to zero, based on their unidentifiability and lack of contribution to improving the model’s fit to the experimental data. For the yeast system, the model reduction process led to significant insights. We found that several reactions initially assumed to be part of the metabolic cycle could be eliminated without compromising the model’s ability to describe the experimental data. Specifically, the pathway from InsP_6_ to 1,5-InsP_8_ via 1-InsP_7_ was not required, and instead, the data were best explained by a linear pathway where 1,5-InsP_8_ is produced directly from 5-InsP_7_. This finding suggests that, in yeast, the conversion of InsP_6_ to 1,5-InsP_8_ primarily occurs through the phosphorylation of 5-InsP_7_, with the dephosphorylation of 1,5-InsP_8_ being the main route for generating 1-InsP_7_.

In the HCT116 cell line, the analysis was slightly more complex due to the presence of two different experimental conditions under which datasets were generated. Both conditions provided valuable insights, but they led to comparable yet slightly different results. The model reduction process revealed a preference for a linear rather than a cyclic metabolic pathway under both conditions. However, there were subtle differences in the parameter estimates and the specific reaction rates. While in the “normal to normal” condition, i.e. under constant P_i_ concentration, only the transitions between 5-InsP_7_ and 1,5-InsP_8_ are removed via the model reduction approach, in the “low to high” phosphate condition, the transition from 1-InsP_7_ to InsP_6_ is additionally removed. Furthermore in the “normal to normal condition, the conversion 1-InsP_7_ to 1,5-InsP_8_ is reduced to a zero order reaction, while in the “high to low” setting it is the other way around. This suggests that the linear pathway might be modulated differently depending on the experimental context. Despite these differences, the overarching conclusion remained consistent: the transition from 5-InsP_7_ to 1,5-InsP_8_ via phosphorylation was not necessary, supporting the notion of a linear reaction sequence in 1,5-InsP_8_ generation in HCT116 cells.

Interestingly, despite these variations, the consistency in the overall pathway structure across different conditions in HCT116 cells, as well as between yeast and HCT116 cells, underscores the robustness of the linear metabolic model. This finding challenges the traditional view of a cyclic IPP metabolic pathway and suggests that the linear pathway might be a more general feature of IPP metabolism in eukaryotic cells.

Our analysis also highlighted the importance of using both experimental data and mathematical modelling to understand complex biochemical networks. The use of pulse-labelling with ^18^O water provided dynamic information on the turnover rates of different IPPs, which was critical for parameter estimation in our ODE models. Additionally, the model reduction approach allowed us to identify the essential reactions in the metabolic pathway, leading to a more parsimonious and interpretable model that still accurately described the experimental data.

In conclusion, this study provides new insights into the metabolic pathways of IPP conversion, revealing that the synthesis of 1,5-InsP_8_ in both yeast and HCT116 cells follows a linear rather than a cyclic pathway. The observation that different experimental conditions in HCT116 cells led to comparable, yet slightly distinct, results suggests that while the linear pathway is robust, its regulation may vary under different physiological contexts. These findings have important implications for our understanding of IPP metabolism and its regulation in eukaryotic cells. Further studies will be needed to explore the functional consequences of this linear pathway and to determine whether similar metabolic networks exist in other organisms or under varying physiological conditions.

## 4 Materials and Methods

### 4.1 Mathematical modelling

The modelling approach presented in this work is based on ODEs where every dephosphorylation, or phosphorylation reaction between the considered InsP species, for all of which experimental data has been provided, was modelled according to Michaelis-Menten kinetics

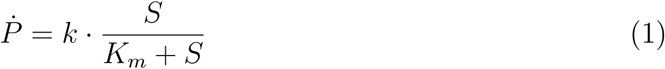

where *P* is the product concentration, *S* is the substrate concentration and *K_M_*is the Michaelis-Menten constant. These can be divided into three regimes, namely: **Zero-order reactions**, where the value of *K_M_* is small compared to *S* and can be neglected, *S* cancels out and the reaction velocity *Ṡ* is independent from the substrate concentration *S*; **Michaelis-Menten reactions**, where *S* is in the range of *K_M_*and the unreduced Michaelis-Menten kinetic has to be applied; **Mass-action reactions**, where *K_M_*is large compared to *S* and the contribution of *S* can thus be neglected in the denominator, meaning the reaction velocity depends linearly on 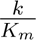 and *S*. *In vitro* the considered dephosphorylation, or phosphorylation reactions can be performed by multiple enzymes, but since in this work only the kinetics of the conversions as a whole are of interest, we consider the sum of the possible enzyme contributions and only model *netto* reactions by a single Michaelis-Menten reaction. To account for the availability of labelled ATP and to describe the labelling of the different molecules, the phosphorylation reaction velocities *k*_i_ are multiplied with the relative abundance of the labelled ATP *ratio*_XO_ where X∈ {0, 1, 2, 3} designates the number of labelled oxygen atoms in the *γ*-phosphate, for which experimental data is also available. In this analysis we considered two cell types, namely baker’s yeast and human HCT116 cells which were both analysed with the identical initial model consisting of 29 states and 48 dynamic parameters. The reactions of the initial full model are shown in Table 1. An absolute error model was assumed, where the parameters of the latter were simultaneously estimated. The observables and the error parameters are shown in Table 2. For all parameters, a logarithmic transformation was performed prior to optimisation to, on the one hand ensure positivity and on the other hand enable optimisation over a wide range of possible values.

**Table 1:**
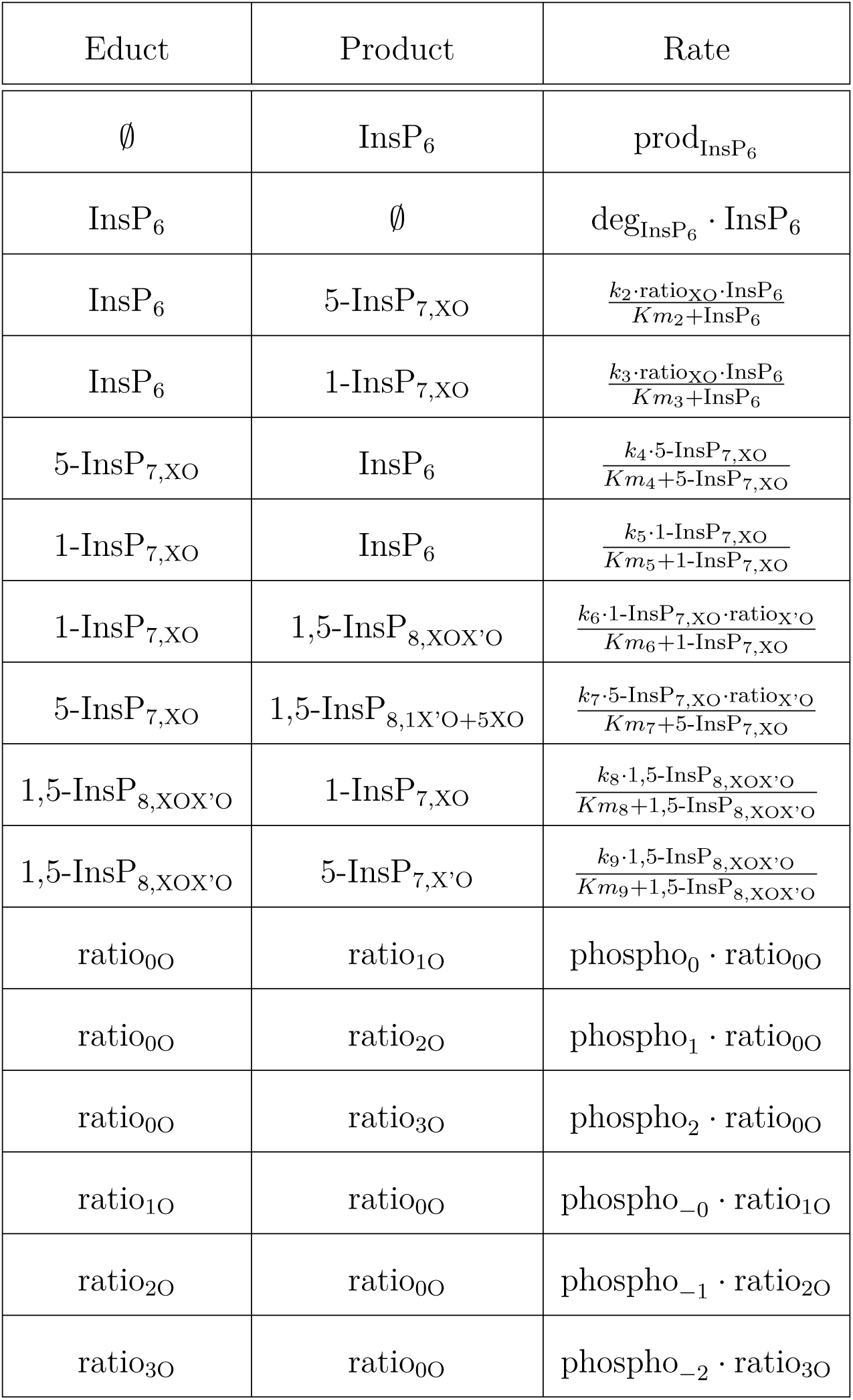
Reactions of the full unreduced mathematical model. The reactions are based on Michaelis-Menten and mass-action kinetics, where X and X’ specify the number of heavy labelled oxygen in the respective phosphate groups, with X and X’∈ {0, 1, 2, 3}.

**Table 2:**
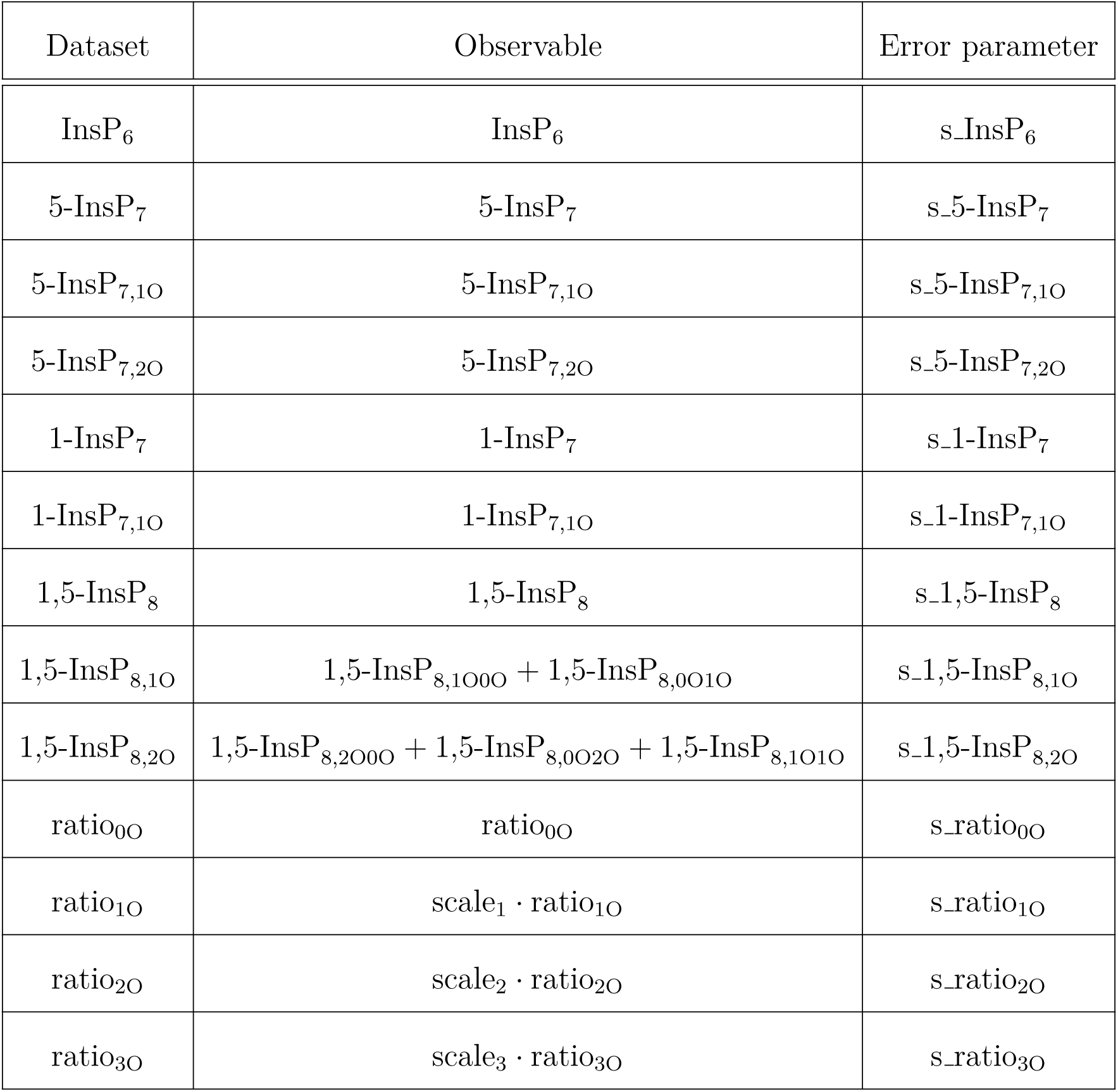
Observables and error parameters of the full unreduced mathematical model.

### 4.2 Parameter estimation and selection of mathematical models

In order to determine the optimal parameter set, the local deterministic Gauss-Newton gradient-based trust-region optimiser was used, which is implemented in the *R* package *dMod* [Kaschek et al., 2019]. To ensure, that the “best” parameter set which is found is also the globally optimal set, a multi-start analyses with 200 random initial parameters sets was performed. To determine how well a fit describes the analysed data the Bayesian information criterion (BIC) is used in this work. The BIC is defined as BIC = *k* ln(*n*) − 2 ln(L), where *k* is the number of model parameters, *n* is the number of data points and L is the maximised value of the likelihood function. Lower BIC values indicate preferable description of the analysed data. Thus, when two models describe the same data comparably well, the model with the lower number of parameters, following Occam’s razor, is to be preferred. It should be mentioned that there exists another popular metric of comparing two models called Akaike information criterion (AIC), defined as AIC = 2*k* − 2 ln(L). However, in this work we use BIC since it is a more stringent criterion and AIC under-penalizes the number of parameters if *n* becomes large. It should be mentioned, that the BIC value is only useful as a relative comparison between different models, while the absolute value of the BIC has no real meaning for any given model.

### 4.3 Profile likelihood

Analysing parameter uncertainties as well as identifiability of the latter is a key task in mathematical modelling. While there are other methods, like Markov-Chain-Monte-Carlo methods (MCMC), to analyse parameter uncertainty, in this project the profile likelihood method was employed [Kreutz et al., 2013, Maiwald et al., 2016]. The profile likelihood is a Likelihood ratio test with one degree of freedom, where the analysed parameter is slightly changed from its best-fit value and all remaining parameters are re-optimised. The ratio between the initial best likelihood value and the new one is then plotted. This process is performed multiple times in a set interval for the analysed parameter, usually +/-3 around the best-fit value on log-scale, and ideally will then give a bell shaped curve passing a 95% confidence level threshold for larger and smaller values of the best-fit value of the parameter. This method cannot only be used to investigate whether model parameters are identifiable, but also provides statistically sound estimates for parameter uncertainties. Furthermore, this method can also guide model reduction by providing not only the parameter uncertainties but also their dependencies on the other parameters in the model. To perform the uncertainty analysis in *R*, the profile likelihood function of the *dMod* package was used.

### 4.4 Yeast strains and genetic manipulation

The Saccharomyces cerevisiae BY4741 was used in this study. The plasmid used to overexpress the truncated Pho90 (Pho90^Δ SPX^) was generated by cloning the Pho90 open reading frame excluding N-terminal 375 amino acids between BamHI and XhoI restriction enzyme sites of the parent plasmid (pRS415) (Mumberg, Muller, and Funk, 1995) containing TEF1 promoter. For DNA transformation, BY4741 strain was logarithmically grown at 30 ^◦^C in 50 mL of YPD (yeast extract-peptone-dextrose) medium. Cultures (4.6 x 10^7^ cells/mL) were centrifuged at 3200 g for 3 min. The pellet was washed with the same volume of TE buffer (10 mM of Tris-HCl pH 7.5 and 1 mM of EDTA) and resuspended with 10 mL of LiAC-TE buffer (TE buffer with 100 mM of LiAC). After the incubation at room temperature for 15 min, cells were collected by centrifugation at 3,000 g for 3 min. The pellet was resuspended with 800 *µ*L of LiAC-TE buffer. 40 *µ*L of cells were mixed with 200 ng of plasmid DNA, 5 *µ*L of salmon sperm DNA (10 mg/mL) and 300 *µ*L of PEG-LiAC-TE buffer (50 % of polyethylene glycol 4000). After 20 min at 30 ^◦^C with gentle shaking, a heat shock was carried out at 42 ^◦^C for 20 min with gentle shaking. Cells were centrifuged at 3,000 g for 2 min, washed twice with water, and grown on selection plates for 2-4 days at 30 ^◦^C. For yeast growth, synthetic complete (SC) medium was prepared from yeast nitrogen base without phosphate (Formedium, UK). The phosphate concentration was adjusted with KH_2_PO_4_. Potassium concentration was controlled by adding KCl instead of KH_2_PO_4_.

### 4.5 Extraction of InsPPs and ATP from yeast cells

Yeast strain overexpressing Pho90^ΔSPX^ was grown at 20 ^◦^C in SC medium containing 0.2 mM of phosphate until logarithmic phase (4.6 x 10^7^ cells/mL). Samples were harvested by centrifugation at 3,000 g for 3 min and resuspended in the same volume of SC medium containing 50 mM of phosphate prepared with 50 % of ^18^O-labelled water. Yeast cells were further incubated at 20 ^◦^C. At each time point, 3 ml of yeast culture was mixed with 300 *µ*l of 11 M perchloric acid to a final concentration of 1 M and snap frozen in liquid nitrogen. InsPPs and ATP were extracted from yeast samples based on [Kim et al., 2024]. Thawed samples were centrifuged at 20,000 g for 3 min at 4 ^◦^C and the soluble supernatant was transferred into a new tube. 6 mg of titanium dioxide (TiO_2_) beads per sample (GL Sciences, Japan) was washed twice with 1 mL of water and 1 M of perchloric acid respectively, and then mixed with the soluble supernatant [Wilson et al., 2015]. The mixture was gently rotated for 15 min at 4 ^◦^C and centrifuged at 20,000 g for 3 min at 4 ^◦^C. After two rounds of washing with 500 *µ*L of 1 M perchloric acid, the TiO_2_ beads were incubated with 300 *µ*L of 3 % (v/v) NH_4_OH at 25 ^◦^C for 5 min with gently shaking to elute InsPPs and ATP. The samples were centrifuged at 20,000 g for 1 min and the eluents were transferred into a new tube. The additional centrifugation was conducted to completely remove the residual TiO_2_ beads from the eluents. The resulting supernatant was dried in a SpeedVac (Labogene, Denmark) at 42 ^◦^C. Samples were kept at -20 ^◦^C until analysis.

### 4.6 Mammalian cell culture and extraction of InsPPs and ATP

InsPPs and ATP data from HCT116 cells under normal-to-normal phosphate (P_i_) conditions were obtained from our previously published work ( [Kim et al., 2024]), while data under low-to-high P_i_ conditions were generated in the present study. The cell culture preparation and InsPP extraction were performed following similar procedures with modifications as described in our previous work. HCT116 cells were cultured for six passages in complete DMEM (Thermo Fisher Scientific, Cat. No. 31966021) supplemented with 10% (v/v) fetal bovine serum (PAN Biotech, Cat. No. P30-3306) and 100 U/mL penicillin-streptomycin (Thermo Fisher Scientific). Cells were grown at 37^◦^C in a humidified incubator with 5% CO_2_ and 98% humidity. The low phosphate medium (0.05 mM P_i_) was prepared using P_i_-free DMEM (Thermo Fisher Scientific, Cat. No. 11971025) supplemented with 100 U/mL penicillin-streptomycin, 1 mM sodium pyruvate, and 0.05 mM phosphate (NaH_2_PO_4_). For high phosphate medium containing 50% ^18^O water, a 2x concentrated medium was prepared from DMEM powder (Sigma, Cat. No. D5648) supplemented with 100 U/mL penicillin-streptomycin, 28.16 mM P_i_ (total 30 mM P_i_, including 1.84 mM P_i_ from the DMEM), and 7.4 g/L sodium bicarbonate. This 2x medium was then diluted with an equal volume of ^18^O-labelled water to yield a final medium containing 50% ^18^O water. For the low-to-high P_i_ experiments, HCT116 cells were seeded in complete DMEM and allowed to adhere for 24 hours. Upon reaching approximately 80% confluency, the cells were cultured in low P_i_ medium (0.05 mM P_i_) for 24 hours, and then shifted to high P_i_ medium (15 mM P_i_) prepared with 50% ^18^O water. At each time point, the medium was removed, and the cells were harvested in 5 mL of 1 M perchloric acid. After snap-freezing in liquid nitrogen, the samples were centrifuged at 3,200 xg for 5 minutes. The soluble supernatant was transferred to a new tube and mixed with TiO_2_ beads (5 mg per sample). InsPPs and ATP were extracted as described previously for yeast cells.

## Acknowledgements

This study was supported by funding by the Deutsche Forschungsgemeinschaft (DFG, German Research Foundation) – Project-ID 499552394 – SFB 1597 and by the state of Baden-Württemberg through bwHPC and the German Research Foundation (DFG) through grant INST 35/159. This study was supported by the Deutsche Forschungsgemeinschaft (DFG) under Germany’s excellence strategy (CIBSS, EXC-2189, Project ID 390939984) H.J. acknowledges funding from the Volkswagen Foundation (VW Momentum Grant 98604). AM is grateful for support by the ERC (Project 788442) and the SNSF (project 10.001.312) and GK for support by HFSPO (LT000588/2019-L).

## A HCT116 low to high phosphate

**Figure 10:**
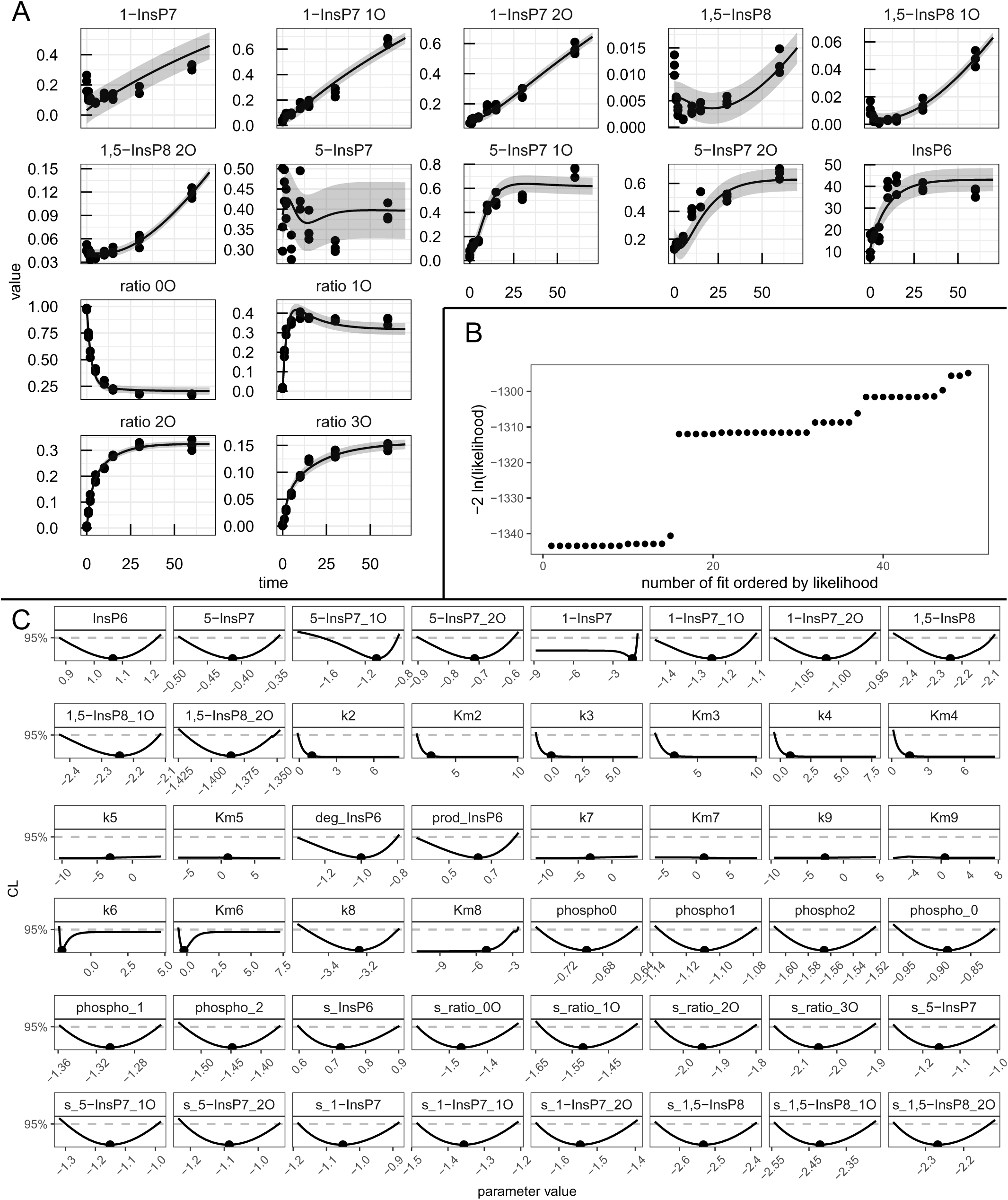
Analysis of the HCT116 data-set with the “full” mathematical model model. (A) Fit of the mathematical model with the best-fit parameters (line) to the experimental data (points). The values are given in *µ*M except for the four species in the lower left of the panel which represent ratios of 1,2,3-labelled to 0-unlabelled *γ*-phosphate. (B) Likelihood values of the 50 best multi-start fits, ordered by lowest likelihood value. (C) Profiles of the best-fit parameters of the full model (line). Best-fit parameter value are represented as points. Parameter values (x-axis) are displayed on log10 scale.

**Figure 11:**
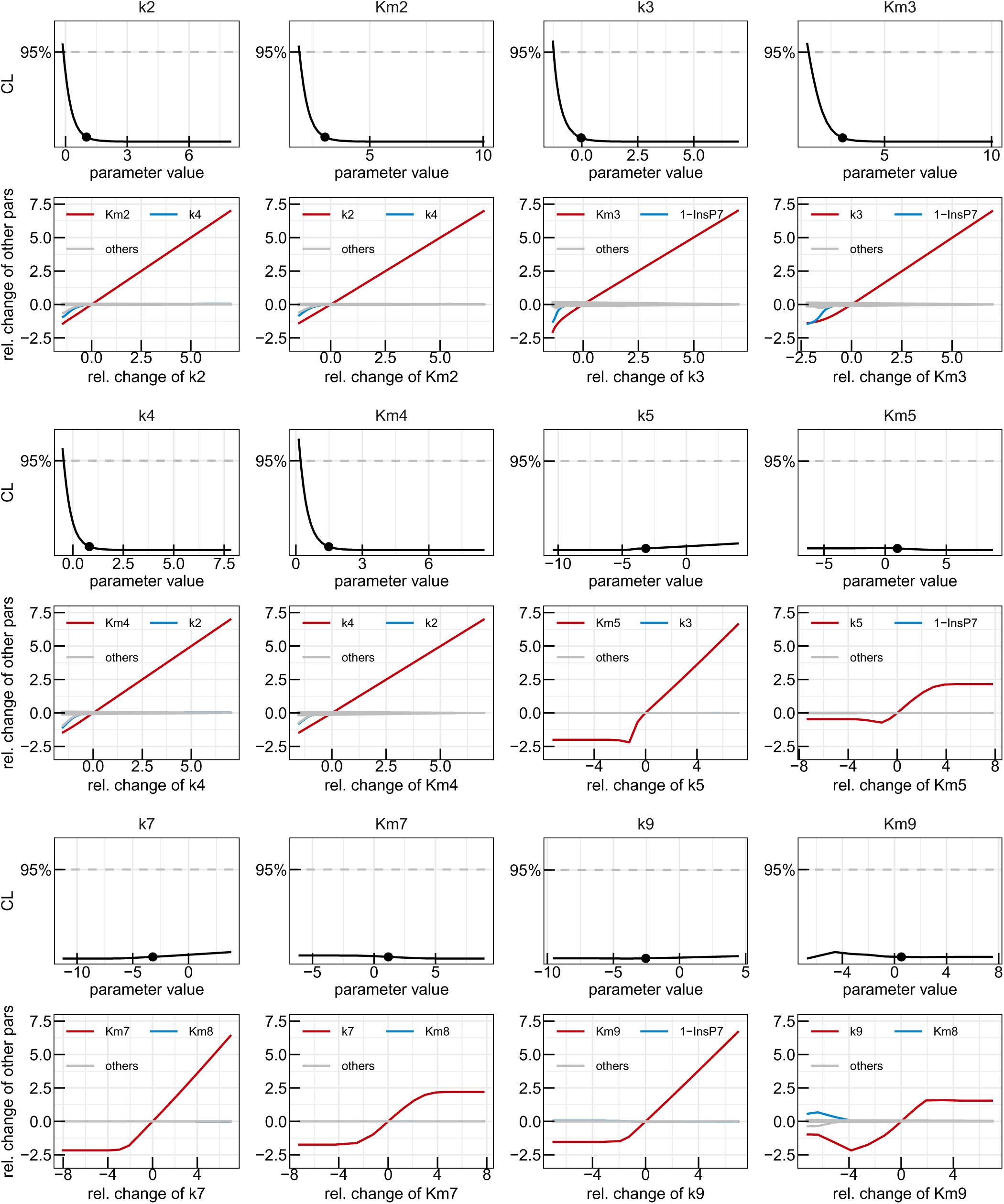
First reduction step in the HCT116 model. Profiles of the parameters considered for the first model reduction step in the yeast metabolic cycle. Best-fit parameter value are represented as points. Parameter values (x-axis) are displayed on log10 scale. Below each profile, the paths visualising the dependencies on the other model parameters are plotted, with the strongest dependency in red, next strongest in blue and the remaining dependencies in grey. Relative change of the profiled parameter (x-axis) and relative change of the other parameters along the profile (y-axis) are displayed on log10 scale.

**Figure 12:**
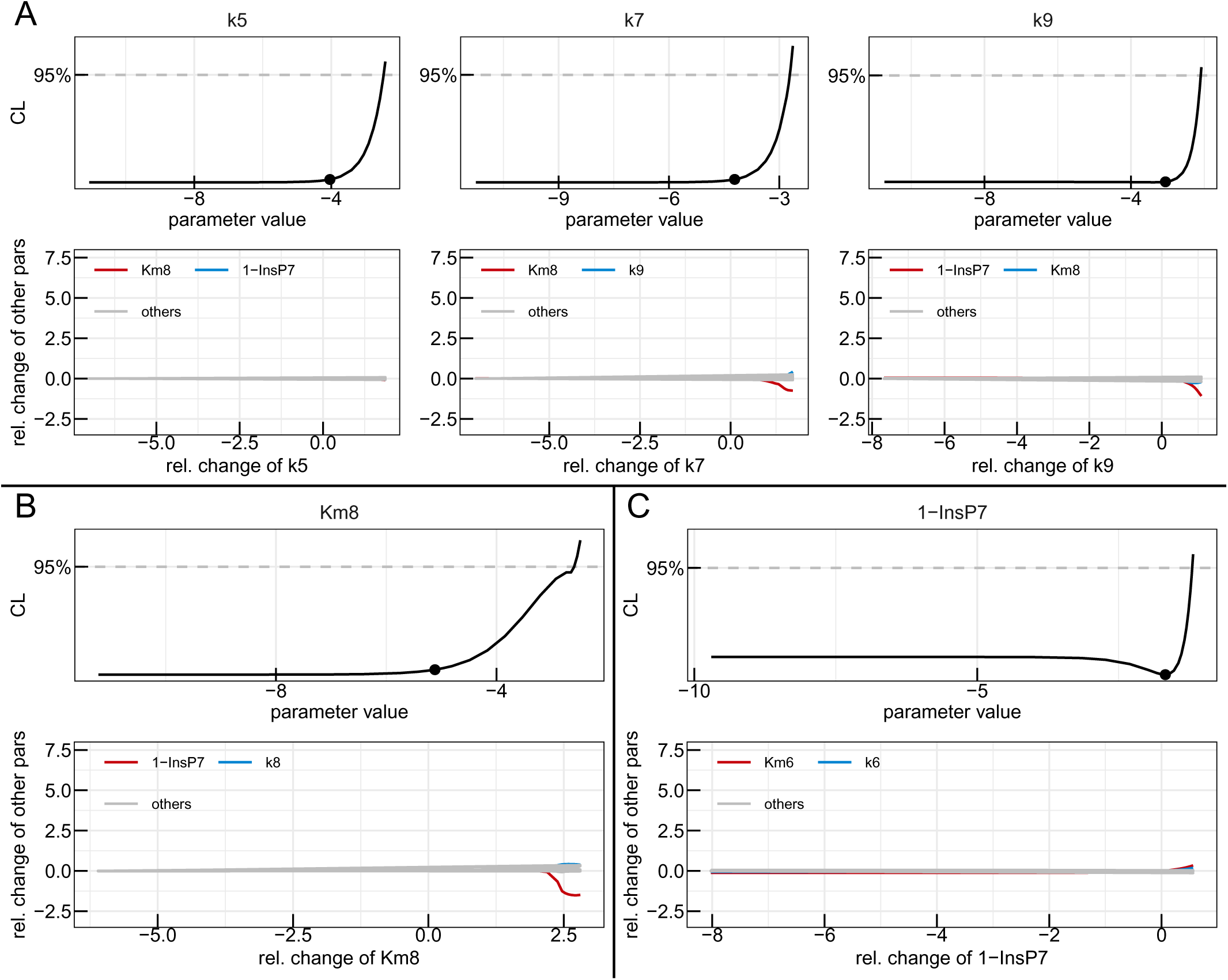
Second and third reduction steps in the HCT116 model. Profiles of the parameters considered for the second (A) and third (B) model reduction step in the yeast metabolic cycle. Best-fit parameter value are represented as points. Parameter values (x-axis) are displayed on log10 scale. Below each profile, the paths visualising the dependencies on the other model parameters are plotted, with the strongest dependency in red, next strongest in blue and the remaining dependencies in grey. Relative change of the profiled parameter (x-axis) and relative change of the other parameters along the profile (y-axis) are displayed on log10 scale.

**Figure 13:**
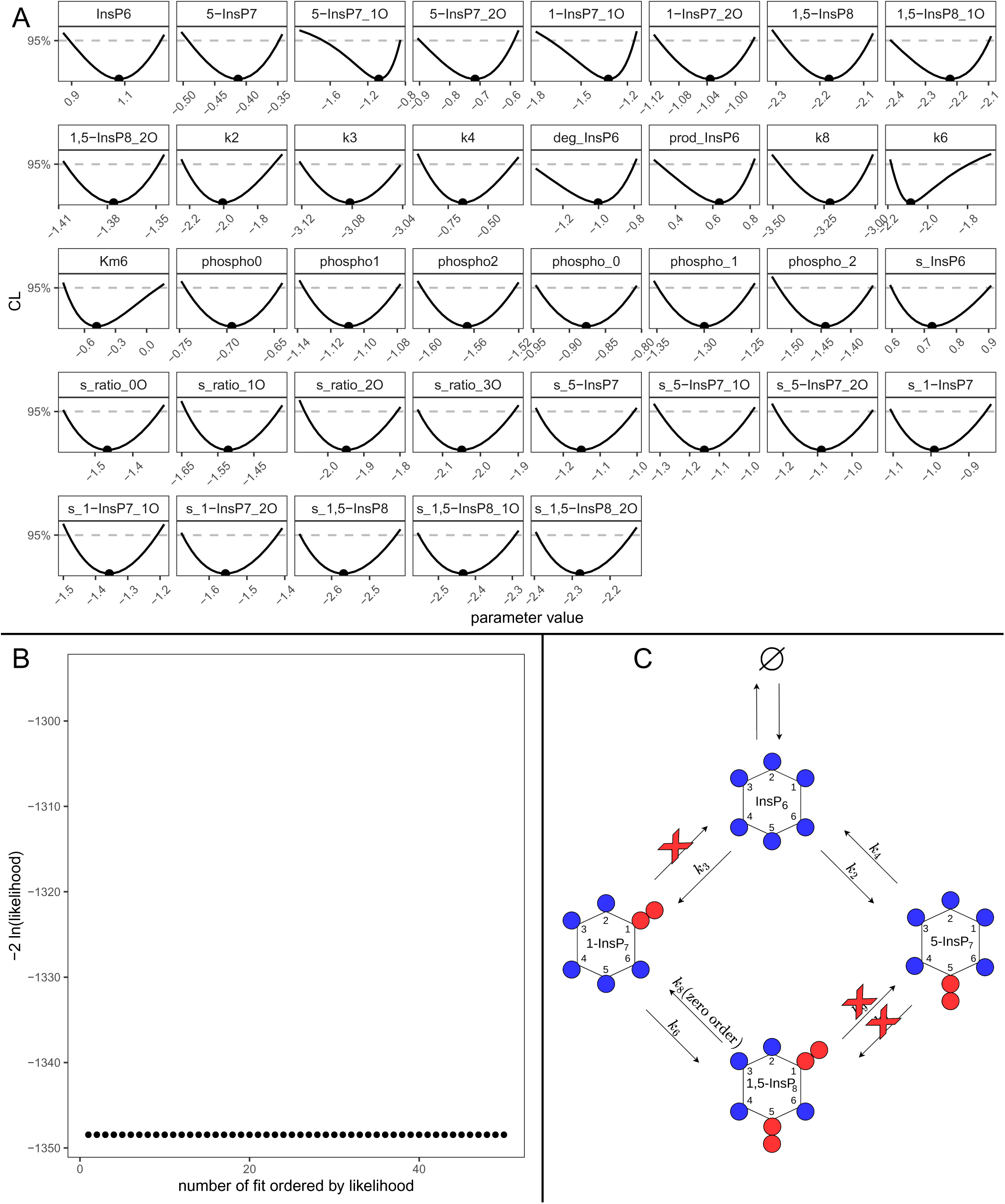
Fully reduced HCT116 model. (A) Profiles of the best-fit parameters of the reduced model (line). Best-fit parameter value are represented as points. Parameter values (x-axis) are displayed on log10 scale. (B) Likelihood values of the 50 best multi-start fits, ordered by lowest likelihood value. (C) Representation of the statistically favoured transition scheme, displaying a chain-like pattern rather than a cycle, with an additional removed transition between 1-InsP_7_ and InsP_6_ compared to the normal-to-normal HCT116 data set.

